# Impact of media brand on cefiderocol disk diffusion results

**DOI:** 10.1101/2024.10.17.618864

**Authors:** MG DeMarco, AM Field, LE Donohue, HL Cox, TS Kidd, KE Barry, AJ Mathers

## Abstract

**Introduction:** Cefiderocol is a siderophore cephalosporin that utilizes iron transport systems to cross cell membranes. This unique strategy complicates antimicrobial susceptibility testing (AST) due to variable iron content in media. While guidance for using iron-depleted media exists for broth microdilution (BMD), disk diffusion (DD) with commercial media is a common AST method in the clinical laboratory. We investigated cefiderocol DD result variability using multiple Mueller-Hinton agar (MHA) brands.

**Methods:** DD results using Remel (Thermo Fisher Scientific, San Diego, CA), Hardy (Hardy Diagnostics, Springboro, OH), and BBL (Becton Dickenson, East Rutherford, NJ) MHA were compared to BMD using iron-depleted cation-adjusted Mueller-Hinton broth. BMD reproducibility, BMD trailing endpoints, and DD intra- and inter-brand variability in zones of inhibition were investigated.

**Results:** Forty-seven multidrug-resistant clinical isolates and three Antibiotic Resistance bank isolates composed of *Pseudomonas aeruginosa*, carbapenemase (CP-) and non-carbapenemase-producing (non-CP-) carbapenem-resistant Enterobacterales (CRE), *Acinetobacter baumannii* complex, *Stenotrophomonas maltophilia*, and *Burkholderia cepacia* complex were tested. Categorical agreement (CA) ≥90% was only demonstrated using CLSI breakpoints with BBL agar. Intra- and inter-brand variability in DD were greatest for *P. aeruginosa* and CRE, with 30% (6/20) and 16.7% (3/18) exhibiting discrepant AST interpretations, respectively. Isolates not susceptible to cefiderocol via BMD were commonly associated with AST interpretation errors and lower CA.

**Conclusions:** Using commercial MHA for DD resulted in frequent AST interpretation discrepancies, particularly for isolates that were not susceptible to cefiderocol by BMD. Methods and quality control may need to be revisited to ensure the reliability of DD for cefiderocol AST.

**IMPORTANCE STATEMENT:** The novel mechanism of action of cefiderocol overcomes a variety of resistance mechanisms associated with gram-negative bacteria and positions the agent as an attractive option for infections involving multidrug-resistant pathogens. The availability of accurate, timely antimicrobial susceptibility testing methods for cefiderocol in clinical microbiology laboratories is critical as cefiderocol-resistant isolates have been described and may contribute to treatment failure. Iron-depleted broth microdilution testing may not be feasible for many clinical laboratories. While disk diffusion is an appealing, practical method to implement, our data demonstrate reproducibility issues across agar brands, most notably for organisms that do not test susceptible to cefiderocol via broth microdilution. Discrepant errors and misclassifications of resistant isolates as susceptible, and susceptible isolates as resistant, may mislead clinicians and compromise treatment efficacy. More work is needed to standardize practical yet reproducible methods for cefiderocol antimicrobial susceptibility testing.

## INTRODUCTION

Cefiderocol is a promising agent for difficult-to-treat and multidrug-resistant (MDR) gram-negative (GN) bacterial infections due to its novel structure composed of a cephalosporin moiety joined to a siderophore. This siderophore provides a unique cell entry method utilizing bacterial iron transport systems to circumvent low outer membrane permeability. It is currently approved by the U.S. Food and Drug Administration (FDA) for use in patients 18 years of age or older for the treatment of complicated urinary tract infections and hospital-acquired or ventilator-associated pneumonia caused by susceptible GN pathogens^1^.

While the mechanism of action of cefiderocol helps overcome some antimicrobial resistance and takes advantage of the low iron environment of infected tissues, demonstrating reproducible antimicrobial susceptibility testing (AST) in an iron-depleted in vitro environment creates challenges for both broth microdilution (BMD) and disk diffusion (DD) modalities^2^. Concerns related to the reproducibility of cefiderocol AST between BMD and DD have been reported^8^, and may relate to a lack of standardized iron content in commercial media^2^. Methods have been established to overcome the impact of iron in BMD by depleting iron from broth used for testing^9–11^, but are likely impractical for many clinical laboratories. Similar guidance does not exist to account for iron content in commercially available Mueller-Hinton agar (MHA) plates used for DD, as iron depletion is not recommended for MHA due to bound iron (approximately 0.5 µg/mL) within the agar resembling an iron-depleted environment^12,13^. However, recent evidence suggests that cefiderocol AST results by DD may vary depending on media brand^14^. Additionally, differences in cefiderocol clinical breakpoints from the Clinical and Laboratory Standards Institute (CLSI)^9^, the European Committee on Antimicrobial Susceptibility Testing (EUCAST)^15^, and the FDA^16^ contribute to discordant susceptibility interpretations. Since clinical failure and treatment-emergent resistance have been described with the use of cefiderocol^3–7^, reliable and accurate AST is critical for guiding effective treatment decisions.

We sought to investigate the impact of MHA brand on the interpretation of DD results for a collection of MDR GN bacterial isolates across multiple clinical breakpoint standards and to assess the concordance of these interpretations with BMD, which served as the reference method.

## METHODS

### Bacterial isolates

Fifty unique GN bacterial isolates including forty-seven de-identified MDR organisms collected from discarded clinical samples between June 2021 and February 2023 (University of Virginia Medical Center Clinical Microbiology Laboratory, Charlottesville, VA) and three Antibiotic Resistance bank isolates (Centers for Disease Control and Prevention, Atlanta, GA) were utlized for BMD and DD testing. Isolates were stored at -80°C.

### Broth microdilution

BMD was performed with plates containing cefiderocol and iron-depleted cation-adjusted Mueller-Hinton broth (ID CAMHB) prepared following CLSI guidance^9,10^. Each isolate was combined with ID CAMHB broth to prepare a consistent inoculum of 0.5 McFarland standard (approximately 10^8^ CFU/mL) via the colony suspension method. Then, each standard was diluted with ID CAMHB to produce a 1:100 dilution to approximate a final concentration of 10^6^ CFU/mL. Fifty µL from each isolate were combined with 50 µL of cefiderocol solutions ranging from 0.063 µg/mL to 128 µg/mL and repeated across 3 rows. Each BMD plate also contained a row of *Escherichia coli* ATCC 25922 as a quality control with in-range minimum inhibitory concentration (MIC), a negative control with ID CAMHB broth only, a growth control (GC) well for each isolate containing no cefiderocol, and a GC well for *E. coli* ATCC 25922 containing no cefiderocol. Plates were incubated at 35°C for 16-20 hours before being read according to CLSI guidance^9,10^. BMD results were accepted for isolates demonstrating adequate growth in the form of a button of ≥ 2 mm or heavy turbidity in each GC well and no more than one skipped well per row. MICs for acceptable organisms with clear endpoints were recorded as the well with the lowest concentration of cefiderocol without growth observed as shown in Figure 1. For organisms with trailing endpoints of faint growth across multiple wells, MICs were recorded as the lowest concentration of cefiderocol where a button of ≤ 1 mm, a light haze, or faint turbidity was observed as shown in Figure 1. Modal MICs for each isolate were calculated and interpreted according to available CLSI, EUCAST, and FDA clinical breakpoints when available for each species (Table 1)^9,15,16^.

**Figure 1.**
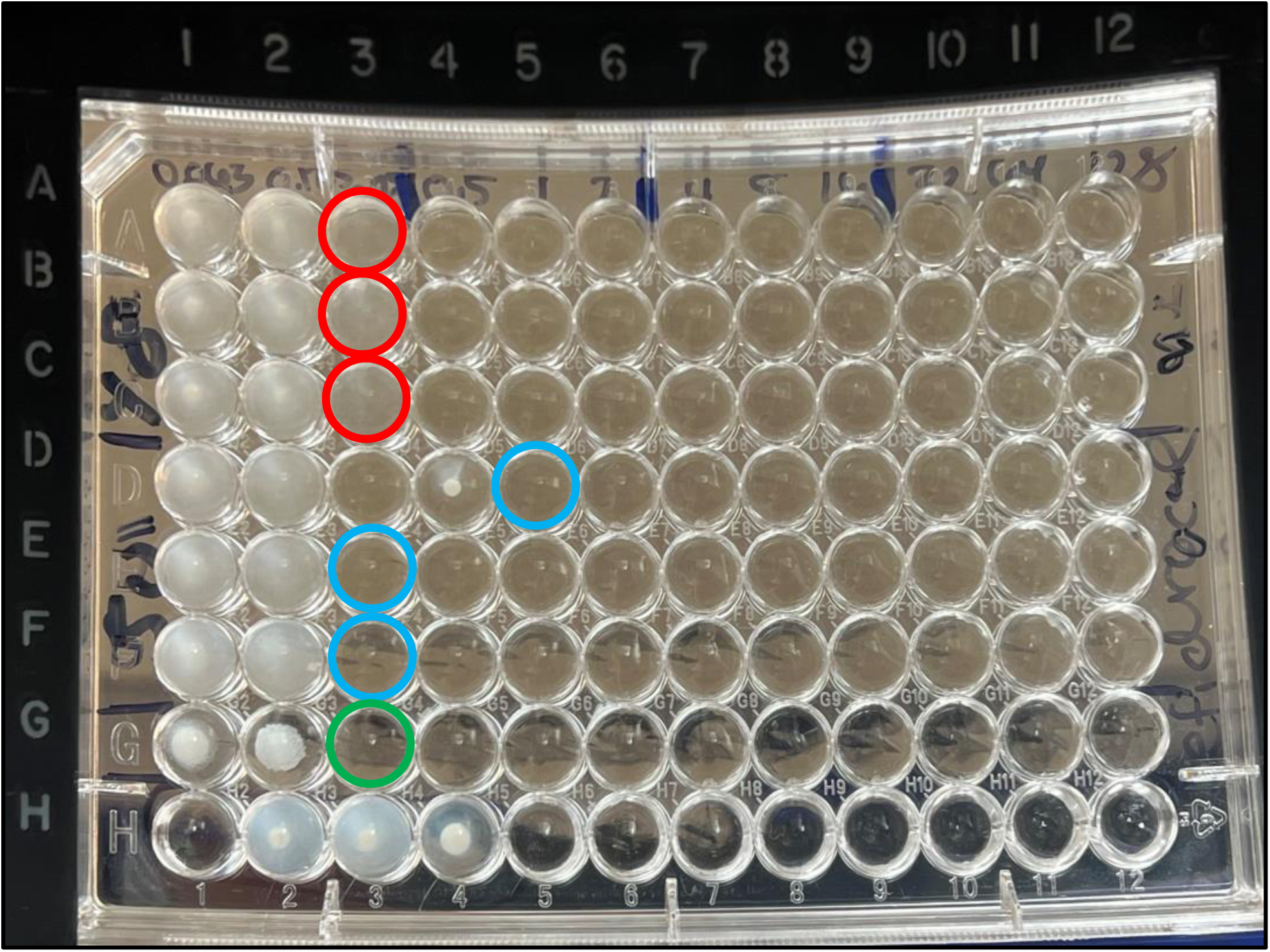
Cefiderocol BMD testing with clear vs. trailing MIC endpoints. Rows A-C = *Stenotrophomonas maltophilia* isolate (Sm2} with hazy trailing endpoints. Red circles denote MICs. Rows D-F = *Stenotrophomonas maltophilia* isolate (Sm4} with clear endpoints. Blue circles denote MICs. Row D featured one skipped well. Row G = *E. coli* ATCC 25922 positive control. Green circle denotes MIC. Row H = Negative control and growth controls for both *Stenotrophomonas maltophilia* isolates and *E. coli* ATCC 25922.

**TABLE 1.** Cefiderocol Susceptibility Testing Interpretive Criteria.

| TABLE 1. Cefiderocol Susceptibility Testing Interpretive Criteria |  |  |  |  |  |  |  |  |
| --- | --- | --- | --- | --- | --- | --- | --- | --- |
| Interpretive Categories and MIC Breakpoints (µg/mL) |  |  |  |  |  |  |  |  |
| Organism | CLSI<sup>a</sup> |  |  | EUCAST<sup>b</sup> |  | FDA<sup>c</sup> |  |  |
|  | S | I | R | S | R | S | I | R |
| Enterobacterales | ≤ 4 | 8 | ≥ 16 | ≤ 2 | > 2 | ≤ 4 | 8 | ≥ 16 |
| *Pseudomonas aeruginosa* | ≤ 4 | 8 | ≥ 16 | ≤ 2 | > 2 | ≤ 1 | 2 | ≥ 4 |
| *Acinetobacter baumannii* complex | ≤ 4 | 8 | ≥ 16 | insufficient evidence |  | ≤ 1 | 2 | ≥ 4 |
| *Stenotrophomonas maltophilia* | ≤ 1 |  |  |  |  | no breakpoints available |  |  |
| *Burkholderia cepacia* complex | no breakpoints available |  |  |  |  |  |  |  |
| Interpretive Categories and Zone Diameter Breakpoints (mm) |  |  |  |  |  |  |  |  |
|  | CLSI<sup>a</sup> |  |  | EUCAST<sup>b</sup> |  | FDA<sup>c</sup> |  |  |
|  | S | I | R | S | R | S | I | R |
| Enterobacterales | ≥ 16 | 9-15 | ≤ 8 | ≥ 23 | < 23 | ≥ 16 | 9-15 | ≤ 8 |
| *Pseudomonas aeruginosa* | ≥ 18 | 13-17 | ≤ 12 | ≥ 22 | < 22 | ≥ 22 | 13-21 | ≤ 12 |
| *Acinetobacter baumannii* complex | ≥ 15 | do not report if ≤ 14 |  | ≥ 17 | insufficient evidence | ≥ 19 | 12-18 | ≤ 11 |
| *Stenotrophomonas maltophilia* | ≥ 15 |  |  | ≥ 20 |  | no breakpoints available |  |  |
| *Burkholderia cepacia* complex | no breakpoints available |  |  |  |  |  |  |  |

<sup>a</sup>Clinical and Laboratory Standards Institute Performance Standards for Antimicrobial Susceptibility Testing, 34th ed.
<sup>b</sup>European Committee on Antimicrobial Susceptibility Testing Breakpoint tables for interpretation of MICs and zone diameters, version 14.0.
<sup>c</sup>United States Food and Drug Administration Identified Breakpoints for Cefiderocol Injection, last updated January 31, 2023.

### Disk diffusion

DD was performed in accordance with CLSI guidance using 30 µg cefiderocol disks (Hardy Diagnostics, Springboro, OH) and three brands of MHA: Remel (Thermo Fisher Scientific, San Diego, CA), Hardy (Hardy Diagnostics, Springboro, OH), and BBL (Becton Dickenson, East Rutherford, NJ)^9,17^. Each isolate was incubated at 37°C on three MHA plates from each brand (for a total of nine plates per isolate) using 0.5 McFarland standard preparations^17^. *Stenotrophomonas maltophilia* isolates were incubated for 20-24 hours, and all others were incubated for 16-18 hours. Mean zones of inhibition for each isolate per plate brand were calculated and interpreted using available CLSI, EUCAST, and FDA clinical breakpoints by species (Table 1)^9,15,16^. Per organism, intra-brand variability was calculated by comparing mean zones of inhibition between the three plates repeated for each brand. Inter-brand variability in terms of range deviation was calculated by comparing mean zones of inhibition across all three plate brands.

### Agreement analysis

Categorical agreement (CA), minor errors (mE), major errors (ME), and very major errors (VME) were assessed by comparing CLSI clinical breakpoint interpretations for mean zones of inhibition determined via DD per brand of MHA to modal MICs determined via BMD as the reference method^9^. Acceptable thresholds for agreement were defined following CLSI guidance, with CA ≥ 90%, mE ≤ 10%, and ME and VME < 3%^18–20^. Acceptable discrepancy rate criteria via the error rate-bounded method were also used in accordance with CLSI guidance^19,20^.

## RESULTS

### Bacterial Isolates

The fifty unique bacterial isolates tested were composed of clinical MDR *Pseudomonas aeruginosa* (n=20), carbapenemase (CP-) and non-carbapenemase-producing (non-CP-) carbapenem-resistant Enterobacterales (CRE) (n=18, n=9 CP-CRE, n=9 non-CP-CRE), *Acinetobacter baumannii* complex (n=6, n=3 from the Antibiotic Resistance bank), *Stenotrophomonas maltophilia* (n=4) and *Burkholderia cepacia* complex (n=2) (Table 2).

**TABLE 2.**
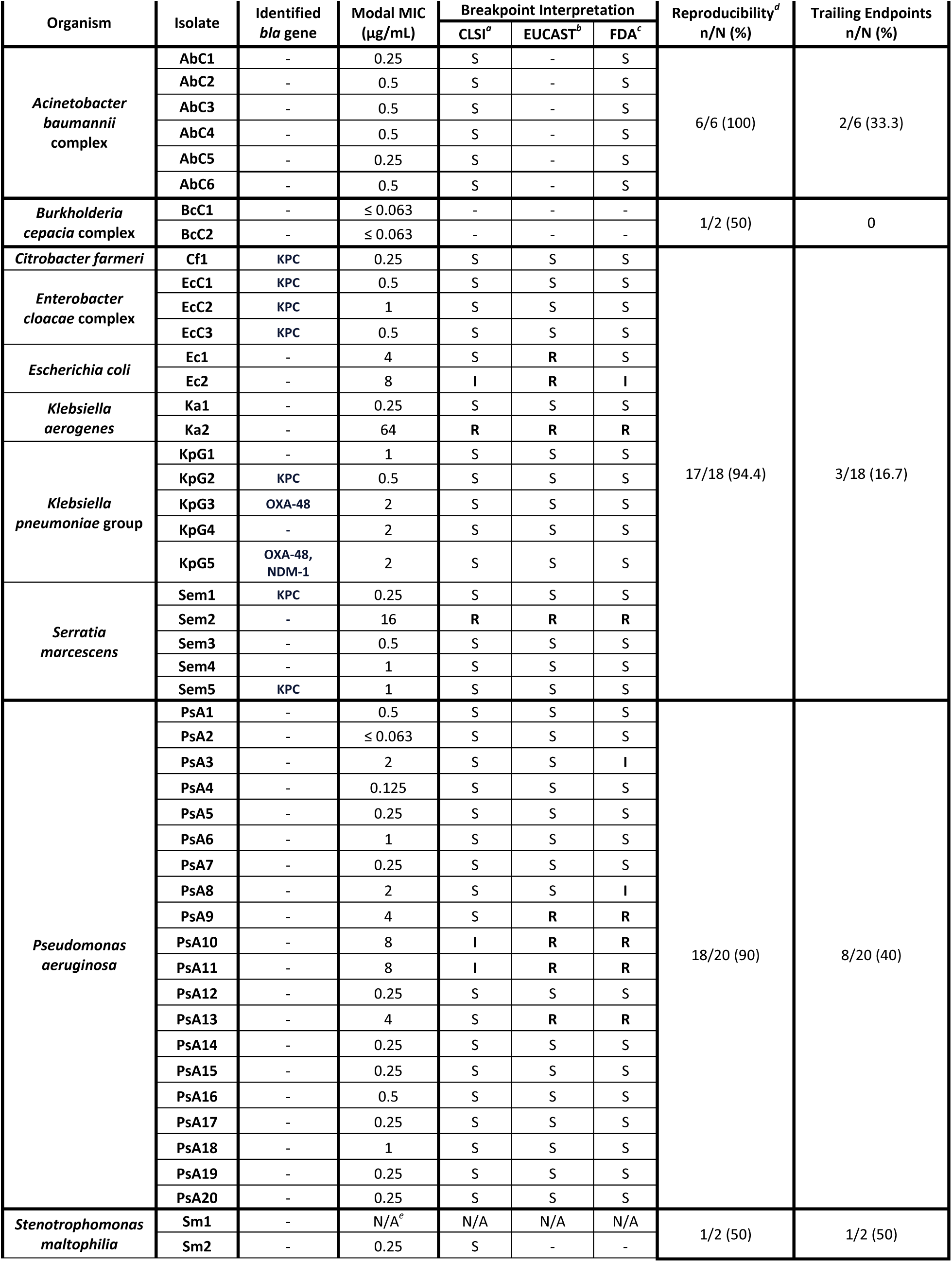
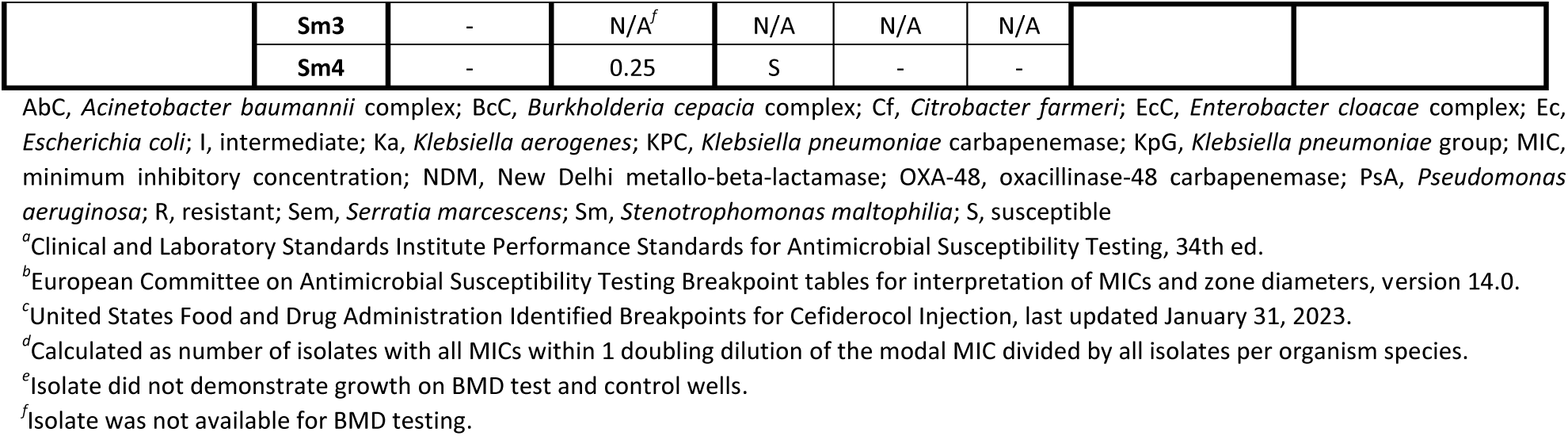
Cefiderocol BMD Results.

### Broth Microdilution

BMD testing was successfully performed on forty-eight isolates (Table 2). Of the forty-six isolates that underwent BMD testing and for which cefiderocol clinical breakpoints exist (i.e., excluding *B. cepacia* complex), most demonstrated in vitro susceptibility, including all *A. baumannii* complex and tested *S. maltophilia* isolates. Intermediate or resistant interpretations per CLSI breakpoints (n=5/46, 10.9%) were recorded for one *Escherichia coli* (Ec2), one *Klebsiella aerogenes* (Ka2), one *Serratia marcescens* (Sem2), and two *P. aeruginosa* (PsA10 and PsA11) isolates. Ten isolates (n=10/46, 22%) demonstrated a modal MIC interpreted as intermediate or resistant by at least one set of established clinical breakpoints (Table 1): two *E. coli* (Ec1 and Ec2), one *K. aerogenes* (Ka2), one *S. marcescens* (Sem2), and six *P. aeruginosa* (PsA3, PsA8, PsA9, PsA10, PsA11, and PsA13). No CP-CRE isolates were interpreted as intermediate or resistant. BMD results for most isolates were within one doubling dilution of their respective modal MICs, demonstrating low variability. Trailing endpoints (Figure 1) were observed for *S. maltophilia* (Sm2; n=1/2; 50%), *P. aeruginosa* (PsA4, PsA8, PsA10, PsA15-17, PsA19, PsA20; n=8/20; 40%), *A. baumannii* complex (AbC2, AbC4; n=2/6; 33%), CP-*C. farmeri* (Cf1; n=1/1; 100%), CP-*E. cloacae* complex (EcC1; n=1/3; 33%), and CP-*K. pneumoniae* group (KpG5; n=1/5; 20%).

### Disk Diffusion

Zones of inhibition and interpretive categorization are outlined in Table 3. Distributions of zones of inhibition for each species and MHA brand are shown in Supplemental Tables 1-15. The average intra-brand variability across replicates was minimal for all species tested: *S. maltophilia* (0.2 mm), *A. baumannii* complex (0.4 mm), CRE (0.6 mm), *P. aeruginosa* (1.1 mm), *B. cepacia* complex (1.2 mm). The average inter-brand variability across media brands was typically greater: *S. maltophilia* (0.6 mm), *A. baumannii* complex (1 mm), CRE (0.9 mm), *P. aeruginosa* (1.8 mm), *B. cepacia* complex (1 mm). For individual isolates, PsA8 demonstrated the largest absolute range (15 mm between Remel and BBL) followed by Ec2 (14 mm between Remel and BBL). Figures 2a and 2b demonstrate noticeable inter-brand variability on examples of *P. aeruginosa* and *E. coli* isolates respectively.

**Figure 2a.**
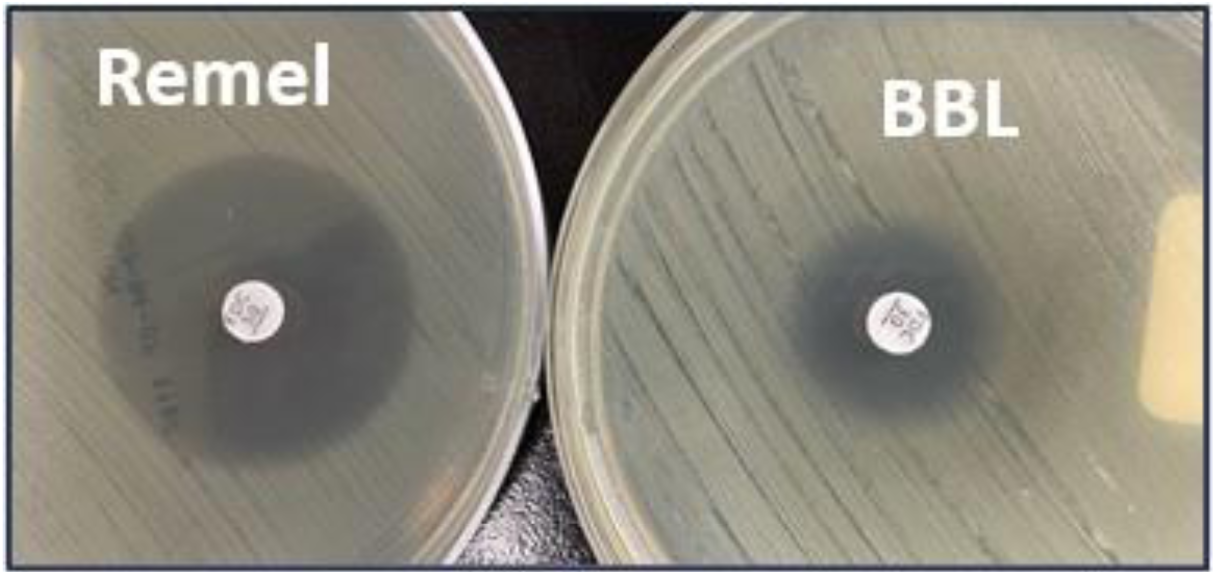
Inter-brand variability in DD zones of inhibition for a *Pseudomonas aeruginosa* isolate (PsA8}. Left: Remel MHA. Right: BBL MHA.

**Figure 2b.**
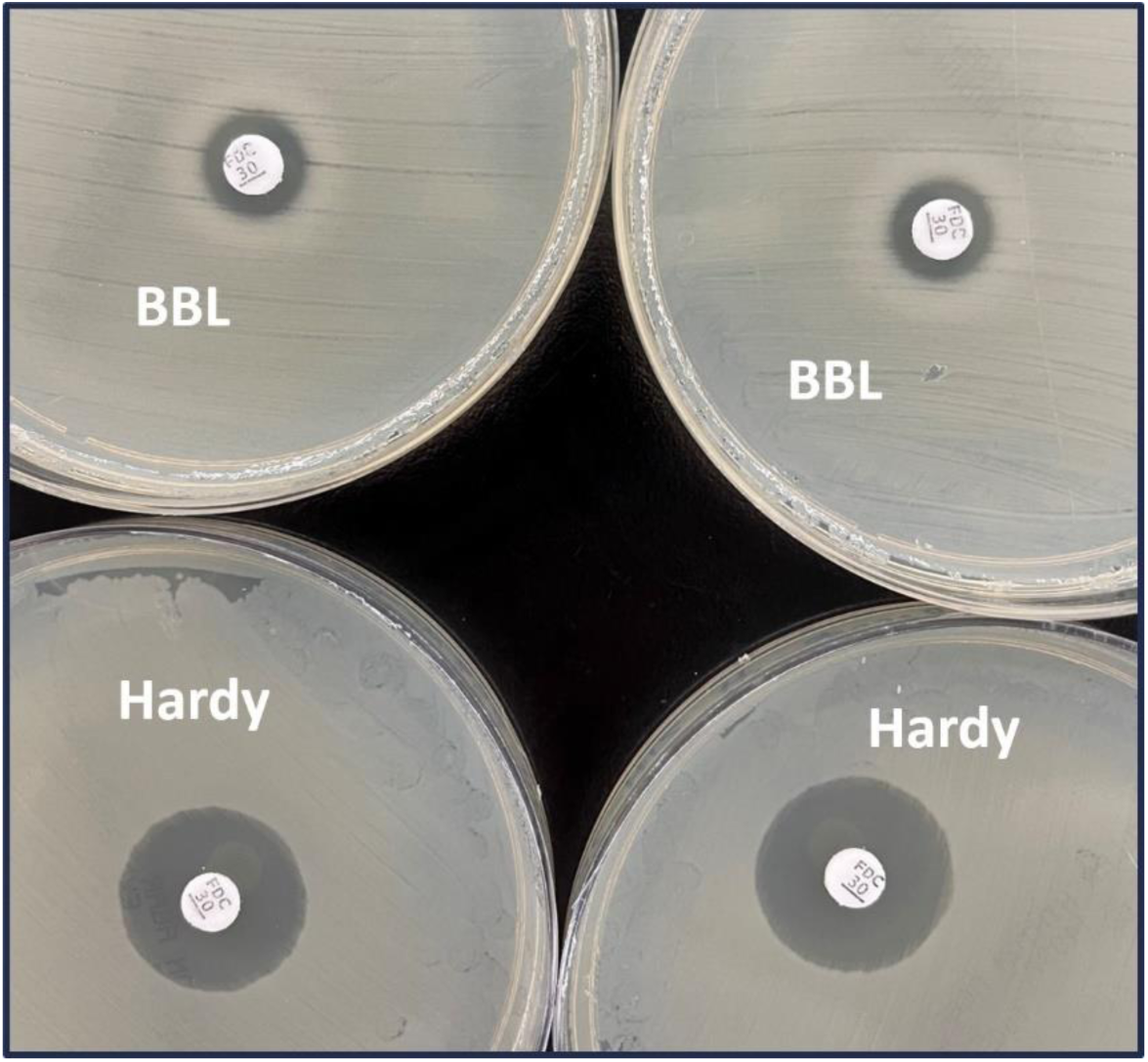
Inter-brand variability in DD zones of inhibition for a carbapenem-resistant *E. coli* (no carbapenemase detected) isolate (Ec2). Top row: BBL MHA. Bottom row: Hardy MHA.

**TABLE 3.** Inter-brand Variability in DD Mean Zones of Inhibition with Cefiderocol Clinical Breakpoint Interpretations.

| Organism | Isolate | Identified *bla* gene | Remel Mean Zone of Inhibition (mm) | CLSI<sup>a</sup> | EUCAST<sup>b</sup> | FDA<sup>c</sup> | Hardy Mean Zone of Inhibition (mm) | CLSI<sup>a</sup> | EUCAST<sup>b</sup> | FDA<sup>c</sup> | BBL Mean Zone of Inhibition (mm) | CLSI<sup>a</sup> | EUCAST<sup>b</sup> | FDA<sup>c</sup> |
| --- | --- | --- | --- | --- | --- | --- | --- | --- | --- | --- | --- | --- | --- | --- |
| *Acinetobacter baumannii* complex | AbC1 | - | 25 | S | S | S | 24 | S | S | S | 24 | S | S | S |
|  | AbC2 | - | 24 | S | S | S | 22 | S | S | S | 23 | S | S | S |
|  | AbC3 | - | 26 | S | S | S | 22 | S | S | S | 21 | S | S | S |
|  | AbC4 | - | 23 | S | S | S | 19 | S | S | S | 21 | S | S | S |
|  | AbC5 | - | 23 | S | S | S | 22 | S | S | S | 21 | S | S | S |
|  | AbC6 | - | 22 | S | S | S | 19 | S | S | S | 20 | S | S | S |
| *Burkholderia cepacia* complex | BcC1 | - | 36 | N/A | N/A | N/A | 36 | N/A | N/A | N/A | 38 | N/A | N/A | N/A |
|  | BcC2 | - | 37 | N/A | N/A | N/A | 35 | N/A | N/A | N/A | 38 | N/A | N/A | N/A |
| *Citrobacter farmeri* | Cf1 | KPC | 26 | S | S | S | 26 | S | S | S | 25 | S | S | S |
| *Enterobacter cloacae* complex | EcC1 | KPC | 25 | S | S | S | 25 | S | S | S | 25 | S | S | S |
|  | EcC2 | KPC | 21 | S | R | S | 21 | S | R | S | 21 | S | R | S |
|  | EcC3 | KPC | 24 | S | S | S | 24 | S | S | S | 23 | S | S | S |
| *Escherichia coli* | Ec1 | - | 23 | S | S | S | 22 | S | R | S | 19 | S | R | S |
|  | Ec2 | - | 24 | S | S | S | 21 | S | R | S | 10 | I | R | I |
| *Klebsiella aerogenes* | Ka1 | - | 28 | S | S | S | 28 | S | S | S | 28 | S | S | S |
|  | Ka2 | - | 10 | I | R | I | 10 | I | R | I | 10 | I | R | I |
| *Klebsiella pneumoniae* group | KpG1 | - | 24 | S | S | S | 24 | S | S | S | 23 | S | S | S |
|  | KpG2 | KPC | 24 | S | S | S | 24 | S | S | S | 25 | S | S | S |
|  | KpG3 | OXA-48 | 24 | S | S | S | 24 | S | S | S | 20 | S | R | S |
|  | KpG4 | - | 26 | S | S | S | 24 | S | S | S | 23 | S | S | S |
|  | KpG5 | OXA-48, NDM-1 | 21 | S | R | S | 20 | S | R | S | 21 | S | R | S |
| *Serratia marcescens* | Sem1 | KPC | 25 | S | S | S | 25 | S | S | S | 24 | S | S | S |
|  | Sem2 | - | 21 | S | R | S | 20 | S | R | S | 20 | S | R | S |
|  | Sem3 | - | 24 | S | S | S | 25 | S | S | S | 24 | S | S | S |
|  | Sem4 | - | 26 | S | S | S | 26 | S | S | S | 24 | S | S | S |
|  | Sem5 | KPC | 30 | S | S | S | 28 | S | S | S | 26 | S | S | S |
| *Pseudomonas aeruginosa* | PsA1 | - | 33 | S | S | S | 30 | S | S | S | 25 | S | S | S |
|  | PsA2 | - | 38 | S | S | S | 37 | S | S | S | 38 | S | S | S |
|  | PsA3 | - | 34 | S | S | S | 32 | S | S | S | 27 | S | S | S |
|  | PsA4 | - | 25 | S | S | S | 26 | S | S | S | 26 | S | S | S |
|  | PsA5 | - | 33 | S | S | S | 33 | S | S | S | 34 | S | S | S |
|  | PsA6 | - | 24 | S | S | S | 24 | S | S | S | 24 | S | S | S |
|  | PsA7 | - | 29 | S | S | S | 27 | S | S | S | 25 | S | S | S |
|  | PsA8 | - | 30 | S | S | S | 25 | S | S | S | 15 | I | R | I |
|  | PsA9 | - | 21 | S | R | I | 20 | S | R | I | 17 | I | R | I |
|  | PsA10 | - | 21 | S | R | I | 20 | S | R | I | 15 | I | R | I |
|  | PsA11 | - | 23 | S | S | S | 22 | S | S | S | 17 | I | R | I |
|  | PsA12 | - | 24 | S | S | S | 24 | S | S | S | 22 | S | S | S |
|  | PsA13 | - | 29 | S | S | S | 29 | S | S | S | 26 | S | S | S |
|  | PsA14 | - | 27 | S | S | S | 27 | S | S | S | 23 | S | S | S |
|  | PsA15 | - | 32 | S | S | S | 31 | S | S | S | 27 | S | S | S |
|  | PsA16 | - | 26 | S | S | S | 27 | S | S | S | 22 | S | S | S |
|  | PsA17 | - | 28 | S | S | S | 31 | S | S | S | 26 | S | S | S |
|  | PsA18 | - | 28 | S | S | S | 27 | S | S | S | 21 | S | R | I |
|  | PsA19 | - | 29 | S | S | S | 26 | S | S | S | 27 | S | S | S |
|  | PsA20 | - | 29 | S | S | S | 21 | S | R | I | 21 | S | R | I |
| *Stenotrophomonas maltophilia* | Sm1 | - | 32 | S | S | N/A | 31 | S | S | N/A | 32 | S | S | N/A |
|  | Sm2 | - | 31 | S | S | N/A | 29 | S | S | N/A | 30 | S | S | N/A |
|  | Sm3 | - | 32 | S | S | N/A | 31 | S | S | N/A | 32 | S | S | N/A |
|  | Sm4 | - | 32 | S | S | N/A | 33 | S | S | N/A | 31 | S | S | N/A |

AbC, *Acinetobacter baumannii* complex; BcC, *Burkholderia cepacia* complex; Cf, *Citrobacter farmeri*; DD, disk diffusion; EcC, *Enterobacter cloacae* complex; Ec, *Escherichia coli*; I, intermediate; Ka, *Klebsiella aerogenes*; KPC, *Klebsiella pneumoniae* carbapenemase; KpG, *Klebsiella pneumoniae* group; NDM, New Delhi metallo-beta-lactamase; OXA-48, oxacillinase-48 carbapenemase; PsA, *Pseudomonas aeruginosa*; R, resistant; Sem, *Serratia marcescens*; Sm, *Stenotrophomonas maltophilia*; S, susceptible
<sup>a</sup>Clinical and Laboratory Standards Institute Performance Standards for Antimicrobial Susceptibility Testing, 34th ed.
<sup>b</sup>European Committee on Antimicrobial Susceptibility Testing Breakpoint tables for interpretation of MICs and zone diameters, version 14.0.
<sup>c</sup>United States Food and Drug Administration Identified Breakpoints for Cefiderocol Injection, last updated January 31, 2023.

**TABLE 3.** KEY.

|  |  |
| --- | --- |
| SUSCEPTIBLE<br>across all brands | Susceptible for selected brand,<br>discrepancy across brands |
| INTERMEDIATE<br>across all brands | Intermediate for selected brand,<br>discrepancy across brands |
| RESISTANT<br>across all brands | Resistant for selected brand,<br>discrepancy across brands |

Of the forty-one isolates that were categorized as susceptible to cefiderocol by BMD via CLSI breakpoints, eight (19.5%) were not susceptible to cefiderocol by DD across MHA brands using CLSI, FDA, and/or EUCAST breakpoints. These discordant isolates included one CP-*E. cloacae* complex (EcC2; n=1/3; 33%), one non-CP-*E. coli* (Ec1; n=1/2; 50%), two CP-*K. pneumoniae* group (KpG3 and KpG5; n=2/5; 40%), and four *P. aeruginosa* (PsA8, PsA9, PsA18, and PsA20; n=4/20; 20%) mostly using EUCAST and FDA breakpoints. Of the *P. aeruginosa* isolates that were susceptible to cefiderocol via BMD with CLSI breakpoints, 2/18 (11%) were not susceptible to cefiderocol by DD testing using BBL MHA and CLSI breakpoints compared to 0/18 when using Remel or Hardy MHA. All fifteen CRE isolates interpreted as susceptible to cefiderocol per CLSI MIC breakpoints (including all CP-CRE) were susceptible per CLSI disk correlates across MHA brands.

Of the five isolates that were categorized as not susceptible to cefiderocol by BMD when applying CLSI breakpoints, four (80%) tested susceptible to cefiderocol by DD across MHA brands and interpretation standards. These discordant isolates included one non-CP-*E. coli* isolate (Ec2) that tested susceptible via DD using Remel and Hardy MHA; one non-CP-*S. marcescens* isolate (Sem2) that tested susceptible using Remel, Hardy, and BBL MHA; and two *P. aeruginosa* isolates (PsA10 and PsA11) that tested susceptible using Remel and Hardy MHA. The only isolate with concordant not susceptible interpretations by both MIC and DD breakpoints was a non-CP *K. aerogenes* (Ka2), adjudicated as resistant per CLSI MIC and EUCAST zone diameter breakpoints, but intermediate per CLSI and FDA disk correlates. Notably, all *A. baumannii* complex and *S. maltophilia* isolates demonstrated susceptibility to cefiderocol by DD across all MHA brands and interpretive criteria, consistent with available BMD results.

### Agreement Analysis

Performance characteristics of DD interpretations compared to BMD as the reference varied across MHA brands when using CLSI, EUCAST, and FDA clinical breakpoints for cefiderocol (Table 1) as shown in Tables 4, 5, and 6, respectively.

**TABLE 4.** Performance characteristics of Remel, Hardy, and BBL MHA DD compared to BMD using CLSI breakpoints*^a^*.

|  |  |  |  |  |  |  |  |  |  |  |  |  |
| --- | --- | --- | --- | --- | --- | --- | --- | --- | --- | --- | --- | --- |
| Organism | CA n/N (%) |  |  | mE n/N (%) |  |  | ME n/N (%) |  |  | VME n/N (%) |  |  |
|  | Remel | Hardy | BBL | Remel | Hardy | BBL | Remel | Hardy | BBL | Remel | Hardy | BBL |
| Overall | 41/46<br>(89.1) | 41/46<br>(89.1) | 42/46<br>(91.3) | 3/46<br>(6.5) | 3/46<br>(6.5) | 3/46<br>(6.5) | 0 | 0 | 0 | 2/46<br>(4.3) | 2/46<br>(4.3) | 1/46<br>(2.2) |
| Susceptible | 41/41<br>(100) | 41/41<br>(100) | 39/42<br>(92.9) | 0 | 0 | 2/41<br>(4.9) | 0 | 0 | 0 | 0 | 0 | 0 |
| Not Susceptible<sup>b</sup> | 0 | 0 | 3/5<br>(60) | 4/5<br>(80) | 4/5<br>(80) | 1/5<br>(20) | 0 | 0 | 0 | 1/5<br>(20) | 1/5<br>(20) | 1/5<br>(20) |
| AbC | 6/6<br>(100) | 6/6<br>(100) | 6/6<br>(100) | 0 | 0 | 0 | 0 | 0 | 0 | 0 | 0 | 0 |
| ≥ I + 2 | N/A | N/A | N/A | N/A | N/A | N/A | N/A | N/A | N/A | N/A | N/A | N/A |
| I+1 to I-1 | N/A | N/A | N/A | N/A | N/A | N/A | N/A | N/A | N/A | N/A | N/A | N/A |
| ≤ I - 2 | 6/6<br>(100) | 6/6<br>(100) | 6/6<br>(100) | 0 | 0 | 0 | 0 | 0 | 0 | 0 | 0 | 0 |
| CRE | 15/18<br>(83.3) | 15/18<br>(83.3) | 16/18<br>(88.9) | 2/18<br>(11.1) | 2/18<br>(11.1) | 1/18<br>(5.6) | 0 | 0 | 0 | 1/18<br>(5.6) | 1/18<br>(5.6) | 1/18<br>(5.6) |
| ≥ I + 2 | 0 | 0 | 0 | 1/1<br>(100) | 1/1<br>(100) | 1/1<br>(100) | 0 | 0 | 0 | 0 | 0 | 0 |
| I+1 to I-1 | 1/3<br>(33.3) | 1/3<br>(33.3) | 2/3<br>(66.7) | 1/3<br>(33.3) | 1/3<br>(33.3) | 0 | 0 | 0 | 0 | 1/3<br>(33.3) | 1/3<br>(33.3) | 1/3<br>(33.3) |
| ≤ I - 2 | 14/14<br>(100) | 14/14<br>(100) | 14/14<br>(100) | 0 | 0 | 0 | 0 | 0 | 0 | 0 | 0 | 0 |
| PsA | 18/20<br>(90) | 18/20<br>(90) | 18/20<br>(90) | 2/20<br>(10) | 2/20<br>(10) | 2/20<br>(10) | 0 | 0 | 0 | 0 | 0 | 0 |
| ≥ I + 2 | N/A | N/A | N/A | N/A | N/A | N/A | N/A | N/A | N/A | N/A | N/A | N/A |
| I+1 to I-1 | 2/4<br>(50) | 2/4<br>(50) | 3/4<br>(75) | 2/4<br>(50) | 2/4<br>(50) | 1/4<br>(25) | 0 | 0 | 0 | 0 | 0 | 0 |
| ≤ I - 2 | 16/16<br>(100) | 16/16<br>(100) | 15/16<br>(93.8) | 0 | 0 | 1/16<br>(6.3) | 0 | 0 | 0 | 0 | 0 | 0 |
| Sm | 2/2<br>(100) | 2/2<br>(100) | 2/2<br>(100) | 0 | 0 | 0 | 0 | 0 | 0 | 0 | 0 | 0 |

AbC, *Acinetobacter baumannii* complex; BMD, broth microdilution; CA, categorical agreement; CRE, Carbapenem-resistant *Enterobacteriales*; DD, disk diffusion; I, intermediate; mE, minor error; ME, major error; MHA, Mueller-Hinton agar; PsA, *Pseudomonas aeruginosa*; R, resistant; Sm, *Stenotrophomonas maltophilia*; VME, very major error. Green shading represents acceptable agreement or error based on CLSI M23 and M52. Red shading represents unacceptable agreement or error based on CLSI M23 and M52.
<sup>a</sup>Clinical and Laboratory Standards Institute Performance Standards for Antimicrobial Susceptibility Testing, 34th ed.
<sup>b</sup>Organisms with intermediate or resistant BMD susceptibility interpretations for cefiderocol according to the 34<sup>th</sup> edition of the CLSI M100.

**TABLE 5.** Performance characteristics of Remel, Hardy, and BBL MHA DD compared to BMD using EUCAST breakpoints*^a^*.

| Organism | CA n/N (%) |  |  | mE n/N (%) |  |  | ME n/N (%) |  |  | VME n/N (%) |  |  |
| --- | --- | --- | --- | --- | --- | --- | --- | --- | --- | --- | --- | --- |
|  | Remel | Hardy | BBL | Remel | Hardy | BBL | Remel | Hardy | BBL | Remel | Hardy | BBL |
| **Overall** | 32/38<br>(84.2) | 32/38<br>(84.2) | 31/38<br>(81.6) | 0 | 0 | 0 | 2/38<br>(5.3) | 3/38<br>(7.9) | 6/38<br>(15.8) | 4/38<br>(10.5) | 3/38<br>(7.9) | 1/38<br>(2.6) |
| **Susceptible** | 28/30<br>(93.3) | 27/30<br>(90) | 24/30<br>(80) | 0 | 0 | 0 | 2/30<br>(6.7) | 3/30<br>(10) | 6/30<br>(20) | 0 | 0 | 0 |
| **Not Susceptible<sup>b</sup>** | 4/8<br>(50) | 5/8<br>(62.5) | 7/8<br>(87.5) | 0 | 0 | 0 | 0 | 0 | 0 | 4/8<br>(50) | 3/8<br>(37.5) | 1/8<br>(12.5) |
| **CRE** | 14/18<br>(77.8) | 15/18<br>(83.3) | 15/18<br>(83.3) | 0 | 0 | 0 | 2/18<br>(11.1) | 2/18<br>(11.1) | 3/18<br>(16.7) | 2/18<br>(11.1) | 1/18<br>(5.6) | 0 |
| **≥ R + 1** | 2/4<br>(50%) | 3/4<br>(75%) | 4/4<br>(100) | 0 | 0 | 0 | 0 | 0 | 0 | 2/4<br>(50%) | 1/4<br>(25%) | 0 |
| **≤ S - 1** | 10/11<br>(90.9) | 10/11<br>(90.9) | 10/11<br>(90.9) | 0 | 0 | 0 | 1/11<br>(9.1) | 1/11<br>(9.1) | 1/11<br>(9.1) | 0 | 0 | 0 |
| **PsA** | 18/20<br>(90) | 17/20<br>(85) | 16/20<br>(80) | 0 | 0 | 0 | 0 | 1/20<br>(5) | 3/20<br>(15) | 2/20<br>(10) | 2/20<br>(10) | 1/20<br>(5) |
| **≥ R + 1** | 2/4<br>(50) | 2/4<br>(50) | 3/4<br>(75) | 0 | 0 | 0 | 0 | 0 | 0 | 2/4<br>(50) | 2/4<br>(50) | 1/4<br>(25) |
| **≤ S - 1** | 14/14<br>(100) | 13/14<br>(92.9) | 12/14<br>(85.7) | 0 | 0 | 0 | 0 | 1/14<br>(7.1) | 2/14<br>(14.3) | 0 | 0 | 0 |

BMD, broth microdilution; CA, categorical agreement; CRE, Carbapenem-resistant *Enterobacterales*; DD, disk diffusion; mE, minor error; ME, major error; MHA, Mueller-Hinton agar; PsA, *Pseudomonas aeruginosa*; R, resistant; VME, very major error. Green shading represents acceptable agreement or error based on CLSI M23 and M52. Red shading represents unacceptable agreement or error based on CLSI M23 and M52.
<sup>a</sup>European Committee on Antimicrobial Susceptibility Testing Breakpoint tables for interpretation of MICs and zone diameters, version 14.0.
<sup>b</sup>Organisms with intermediate or resistant BMD susceptibility interpretations for cefiderocol according to version 14.0 of the EUCAST Breakpoint tables for interpretation of MICs and zone diameters.

**TABLE 6.**
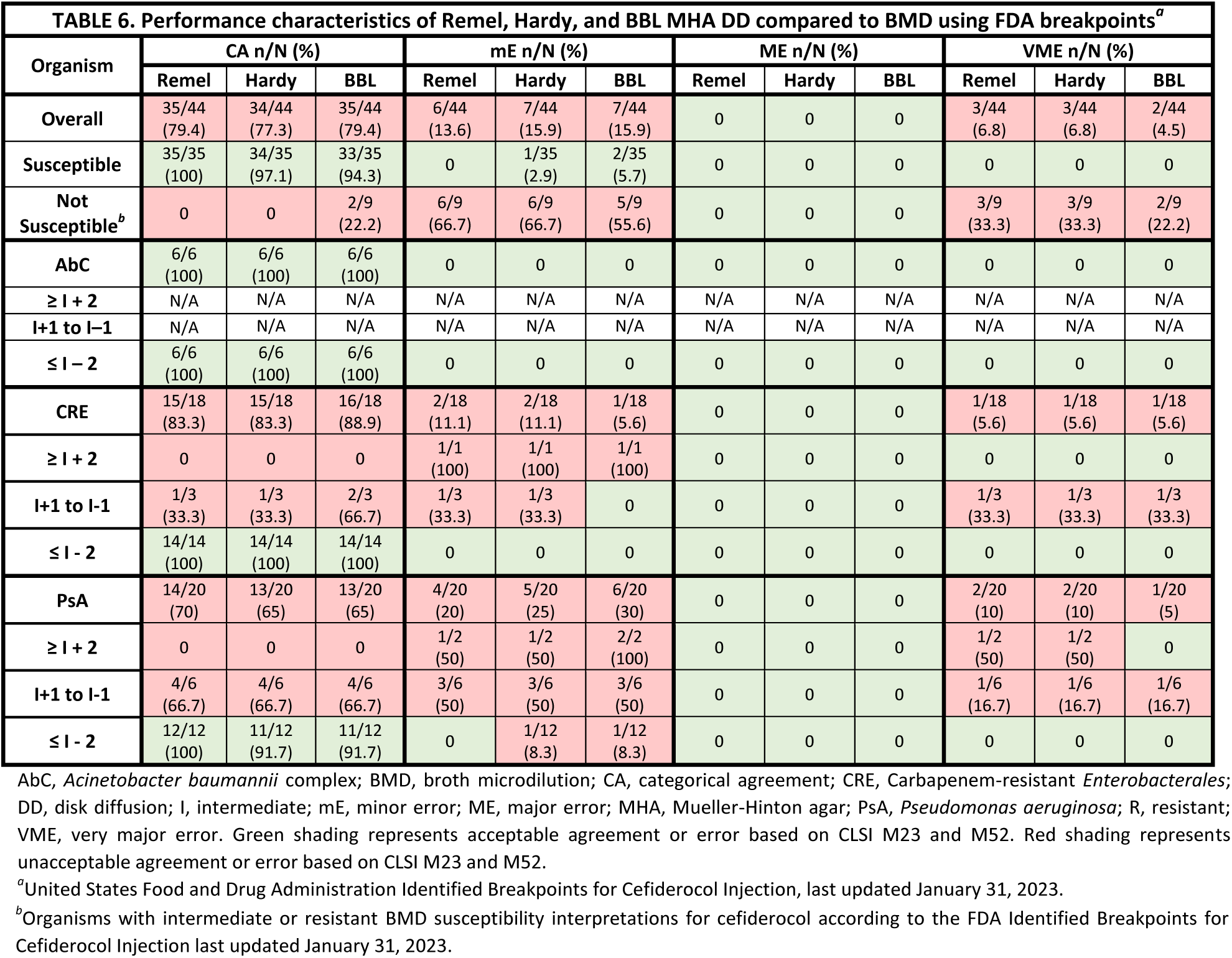
Performance characteristics of Remel, Hardy, and BBL MHA DD compared to BMD using FDA breakpoints*^a^*.

### CLSI Breakpoints

Among forty-one isolates that were susceptible to cefiderocol by BMD, CA was acceptable for all MHA brands (Table 4): Remel (41/41), Hardy (41/41), and BBL (39/41, 95%), with 2 mEs among *P. aeruginosa* isolates (PsA8 and PsA9). However, acceptable CA was not achieved for five non-susceptible isolates via BMD when compared to DD using any MHA brand: Remel (0/5), Hardy (0/5), BBL (3/5, 60%). Considering errors for all CRE isolates regardless of BMD AST interpretation, CA was not acceptable using any MHA (Remel 15/18, 83%; Hardy 15/18, 83%; BBL 16/18, 89%). Acceptable CA was achieved among *P. aeruginosa* isolates (Remel, Hardy, and BBL all 18/20, 90%).

### EUCAST Breakpoints

Thirty isolates were considered susceptible to cefiderocol by BMD using EUCAST breakpoints (Table 5). Acceptable CA was achieved when using Remel (28/30, 93%) and Hardy (27/30, 90%) MHA, but not BBL (24/30, 80%). No mEs or VMEs were observed, but MEs were common. The eight isolates that were not susceptible by BMD fared worse, with CA observed in only 4/8 (50%) isolates with Remel MHA, 5/8 (63%) with Hardy MHA, and 7/8 (88%) with BBL MHA. This was driven by VMEs across all MHA brands. Considering errors among all CRE isolates, acceptable CA was not achieved using Remel (14/18, 78%), Hardy (15/18, 83%), or BBL MHA (15/18, 83%) due to MEs and VMEs. Among all *P. aeruginosa* isolates, Remel MHA yielded an acceptable CA (18/20, 90%), but Hardy (17/20, 85%) and BBL (16/20, 80%) did not due to MEs and VMEs.

### FDA Breakpoints

Among thirty-five isolates that were susceptible to cefiderocol by BMD using FDA interpretations, acceptable CA was achieved across all MHA brands (Table 6): Remel 35/35; Hardy 34/35, 97%; BBL 33/35, 94%. The nine isolates that were not susceptible by BMD again fared worse due to mEs and VMEs. Acceptable CA was not achieved between BMD and DD using Remel or Hardy MHA, and only two of these isolates (2/9, 22%) demonstrated CA using BBL MHA. Acceptable CA was not achieved using any MHA brand among CRE (Remel 15/18, 83%; Hardy 15/18, 83%; or BBL 16/18, 89%) or *P. aeruginosa* (Remel 14/20, 70%; Hardy 13/20, 65%; BBL 13/20, 65%) isolates. All *A. baumannii* isolates demonstrated acceptable CA with no errors observed across MHA brands.

## DISCUSSION

Identifying a testing method that is both practical to implement and capable producing accurate results will reinforce the clinical utility of cefiderocol for the effective treatment of MDR GN bacterial infections. However, among our collection of MDR clinical isolates and Antibiotic Resistance bank isolates, the use of commercial Remel, Hardy, and BBL MHA for DD frequently resulted in AST interpretation discrepancies when compared to BMD. Isolates that were not susceptible to cefiderocol via BMD commonly tested susceptible via DD across MHA brands and interpretation standards, posing a risk of under-calling resistance when using DD for organisms with MICs approaching or exceeding clinical breakpoints via BMD. While our agreement analyses were poorest when only including isolates that were classified as not susceptible to cefiderocol via BMD, analyses that included all tested isolates with available clinical breakpoints still failed to consistently achieve acceptable CA of ≥90% between DD and reference BMD results across MHA brands and breakpoint interpretations. A potential exception to this trend involved CLSI interpretations applied to DD using BBL MHA, but even BBL MHA performed inadequately with isolates that were not susceptible to cefiderocol by BMD. Perhaps most concerning for clinical practice was the observance of VME in which the DD method failed to identify resistance in >3% of isolates across brands and breakpoints with the exception of BBL MHA plates with CLSI (2.2%) and EUCAST (2.6%) breakpoints. These findings align with previous analyses^8,21–23^. Morris et al. proposed that DD may be a convenient alternative to BMD for cefiderocol AST, but observed inconsistencies in susceptibility interpretations across CLSI, EUCAST, and FDA breakpoints for GN bacteria^21^. Bonnin et al. raised further concern regarding DD reliability in a large study of CRE isolates plated on Biorad (Marnes la Coquette, France) MHA^8^. A CA rate of 81.7% and a VME rate of 23.3% prompted a recommendation for confirmatory BMD testing in many isolates to avoid under-calling resistance^8^. Additionally, Liu et al. evaluated the correlation between cefiderocol AST DD and BMD results for a collection of 468 *A. baumannii* complex isolates and found that DD was unable to accurately classify all 6 isolates with cefiderocol MICs ≥ 8 µg/mL as resistant^22^. Jeannot et al. published similar findings with 97 *Acinetobacter* spp. strains and Mast Diagnostic (Merseyside, United Kingdom), Liofilchem (Roseto degli Abruzzi, Italy), and Oxoid (Thermo Fisher Scientific, Basingstoke, United Kingdom) cefiderocol 30µg disks plus Becton Dickinson MHA and suggested that while DD may be a helpful AST screening tool for cefiderocol-susceptible strains, interpretations of strains with zones of inhibition ≤ 22 mm via DD should be confirmed with BMD^23^. Others have echoed this suggestion for BMD confirmatory testing when DD results fall within areas of technical uncertainty for a variety of GN bacteria^9,24,25^. The CLSI M100 also includes a related cautionary comment outlining the potential for inaccurate testing and interpretation^9^. Combined with our analyses, these findings highlight concerns with the use of DD in the pursuit of the identification of reliable yet practical AST methods for cefiderocol especially to identify resistant isolates.

In addition to discrepancies between DD and BMD, DD variability by MHA brand alone could impede accurate and reproducible cefiderocol DD between laboratories. For the species included in our study, this was most worrisome among *P. aeruginosa* and CRE isolates; more than 25% and almost 20% of these isolates demonstrated at least one DD breakpoint interpretation dependent on MHA brand, respectively. Inter-brand variation in zones of diffusion >10 mm was also most common among isolates from these species. No differences in interpretation were noted for *S. maltophilia* or *A. baumannii* complex isolates, but these were all cefiderocol susceptible via BMD, which again raises suspicion that DD cannot reliably differentiate resistant isolates across media. Similarly, while clinical breakpoints do not exist for *B. cepacia* complex, we observed very low modal MICs and large mean zone diameters that were similar across MHA brands. Potter et al. previously investigated the impact of MHA brand on DD interpretation and found similar discrepancies in zones of inhibition between Hardy (Santa Monica, CA) and BD (Franklin Lakes, NJ) MHA that resulted in interpretation differences for 8.7% (2/23) of their *P. aeruginosa* isolates using CLSI breakpoints^14^.

Inconsistent AST results between MHA brands may be a consequence of the interplay between a lack of iron standardization in MHA plates and the unique cell entry method of cefiderocol as a siderophore cephalosporin^2^. The absence of cefiderocol resistance observed among our CP-CRE isolates may be explained by the likelihood that its development depends on the combined influence of multiple mechanisms beyond the expression of carbapenemases including, but not limited to, alterations to siderophore receptors, penicillin-binding-protein-3 target modifications, and loss of function mutations in the *cirA* iron transporter gene^3^. The utilization of iron channels offers a novel strategy for overcoming antimicrobial resistance in GN bacterial infections with limited treatment options^1^. However, it also creates in vitro AST challenges under conditions that do not closely mimic the low iron environment of infected tissues. Variability in BMD results related to iron content prompted warnings from both CLSI^2^ and EUCAST^26^ and led to the subsequent development of methods for depleting iron from broth for testing^9–11^. Similar methods do not exist for commercial DD MHA, which was thought to resemble an iron-depleted environment due to the bound nature of its iron content^12,13^. This may not be accurate and it may be important to develop quality controls or standard media to account for iron content in commercial MHA.

Beyond reproducibility concerns, inconsistencies in cefiderocol DD results across MHA brands may impact the ability to estimate in vivo clinical efficacy from AST results, ultimately impacting clinical decision-making and outcomes. While surveillance studies have shown promising in vitro susceptibility data for cefiderocol against a variety of GN pathogens^27^, clinical data have highlighted concerning trends in clinical response, mortality rates, and the emergence of resistance.^3–7^. For example, Hoellinger et al. described a clinical response rate of 20% and a 30-day mortality rate of 60% for infection-related causes in a small study of mostly immunocompromised patients who received cefiderocol for at least 48 hours for the treatment of a variety of infections attributed to MDR non-fermenting, GN bacilli^4^. Unreliable cefiderocol AST methods further complicate these trends, and impractical methods can contribute to delays in therapy for those with limited options for effective antimicrobials on account of multidrug resistance.

Our study was not without limitations. The trend of discrepancies being highest in the setting of cefiderocol resistance was observed among *P. aeruginosa* isolates and non-CP-CRE isolates in our study, but no other isolates were adjudicated as being intermediate or resistant among other tested species. Although this is true cefidericol resistance remains relatively rare and thus we feel our data set still has value in highlighting a potential issue of variability by media brand. Trailing endpoints were observed during the BMD testing phase of this study for every organism category except *B. cepacia* complex. Although CLSI^9^ and EUCAST^28^ have described strategies for interpreting BMD results despite this phenomenon, there remain opportunities for intra- and inter-reader variability. Despite these limitations, our results demonstrated key inconsistencies in susceptibility interpretations and CA across three MHA brands and breakpoint criteria that warrant further study to identify an accurate yet practical strategy for conducting cefiderocol AST.

## CONCLUSIONS

There is a significant clinical need for standardizing cefiderocol susceptibility testing such that methods are accurate and reproducible. In our study, DD AST for cefiderocol frequently demonstrated interpretation discrepancies across clinical breakpoint standards and commercially available MHA when compared to BMD. Errors, including those that under-called resistance, were most common among isolates that were not susceptible to cefiderocol by BMD, *P. aeruginosa* isolates, and non-CP-CRE isolates. Our results warrant caution with susceptibility interpretation via DD with commercially available MHA when cefiderocol is indicated. Future studies are needed to identify accurate yet practical methods for performing cefiderocol AST and to determine which unique media-related DD results most accurately reflect in vivo efficacy.

## ACKNOWLEDGEMENTS

University of Virginia Medical Center Clinical Microbiology Laboratory, Charlottesville, VA

